# A Natural Programmable Metamaterial Controls 3D Curvature of Compound Eyes

**DOI:** 10.1101/2025.11.25.689431

**Authors:** Juan Garrido-García, Rhian F. Walther, Jesús Torres-Tirado, Jesús A. Andrés-San Román, José A. Sanz-Herrera, Franck Pichaud, Fernando Casares, Luis M. Escudero

**Affiliations:** Departamento de Matemática Aplicada I, Universidad de Sevilla and Instituto de Biomedicina de Sevilla (IBiS), Hospital Universitario Virgen del Rocío/CSIC/Universidad de Sevilla. 41013 Seville, Spain; Cell Biology of Tissue Architecture and Physiology Group. Laboratory for Molecular Cell Biology (LMCB), University College London, London WC1E 6BT, United Kingdom; CABD, CSIC/Universidad Pablo de Olavide/Junta de Andalucía, 41013 Seville, Spain; Departamento de Biología Celular, Facultad de Biología, Universidad de Sevilla and Instituto de Biomedicina de Sevilla (IBiS), Hospital Universitario Virgen del Rocío/CSIC/Universidad de Sevilla. 41013 Seville, Spain; Escuela Técnica Superior de Ingeniería. Universidad de Sevilla, Seville 41092, Spain and Instituto de Biomedicina de Sevilla (IBiS), Hospital Universitario Virgen del Rocío/CSIC/Universidad de Sevilla, 41013 Seville, Spain; Institute for the Physics of Living Systems, University College London, London WC1E 6BT, UK

## Abstract

The panoramic vision of the convex compound eyes, common to insects and crustaceans, relies on micrometer-scale curvature variations^1^. These variations create specialized visual zones adapted to specific tasks, including detecting prey, mates, or predators^2,3^. However, the mechanisms by which such fine-scale curvature is encoded during development remain unknown. Here we show that the developing eye of *Drosophila melanogaster* functions as a natural metamaterial that programs the organ’s precise 3D curvature. We discover a supracellular triangular mesh in the basal retina with a specific pattern of triangles sizes. Computational simulations demonstrate its role directing the small scale curvature variations of the eye. Genetic disruption of this micropattern prevents local curvature establishment. Furthermore, the presence of a homologous mesh-curvature relationship in *Drosophila mauritiana* indicates evolutionary conservation of this mechanism. These results reveal a novel mechanism of morphogenesis control in which the supracellular 2D patterning give rise to a biological programmable metamaterial that encodes 3D curvature with great precision^4^. Our *in vivo* finding offers a novel framework for the design of shape-programmable 3D biological surfaces with broad implications from synthetic morphogenesis to clinical applications.

## Main

Morphogenesis, the controlled shaping of living materials, is essential for the correct organization and function of complex organs. A paradigmatic example of how form impacts function, is the insect compound eye, an optical device of great precision. It consists of a crystalline packing of unit eyes, called ommatidia, into a convex, dome-like structure (Fig. 1a,b). Each ommatidium comprises a central photodetection cartridge capped by a facet lens and ensheathed by a layer of ancillary cells (Fig. 1c). Importantly, curvature can vary across the eye, and this curvature anisotropies often differ between species^1^. Curvature variation modifies visual performance: zones of low curvature focus many ommatidia onto a narrow region, resulting in high spatial-resolution vision, while those with high curvature expand the field of view. The combination of curvature and lens diameter, with larger lenses providing greater light sensitivity, gives compound eyes multiple optical properties. These specialized regions support predation, mating, or escape responses, which are all vital to the animal^2,3^ (Fig. 1d). Therefore, there must be mechanisms responsible for controlling the local curvature of compound eyes, as this trait critically impacts their visual performance.

**Fig. 1.**
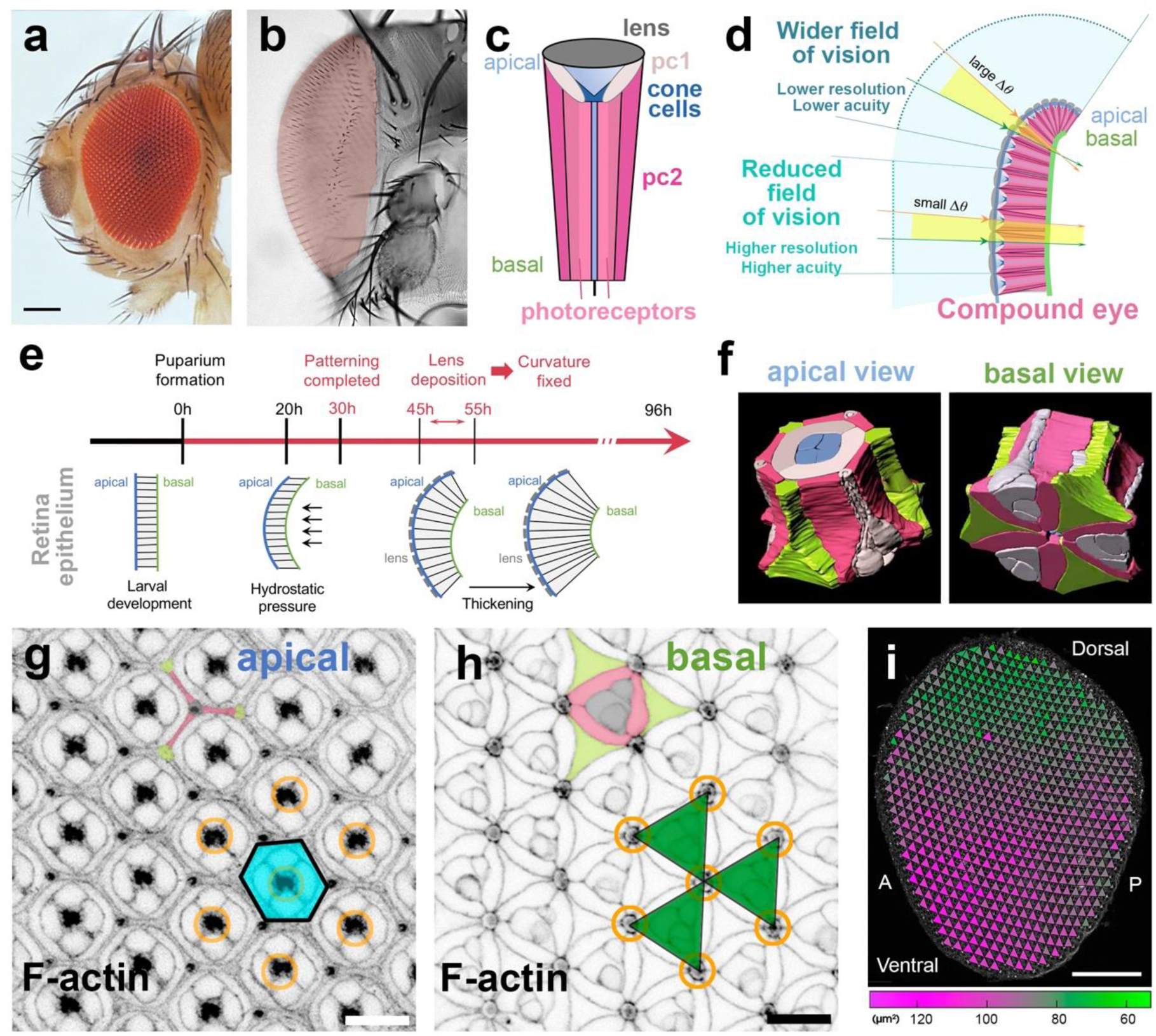
A supracellular triangular mesh patterns the basal pupal retina of *Drosophila*. (**a** and **b**) *D. melanogaster* compound eye. Lateral (a) and frontal (b) views (b, eye pseudocolored in light pink). (**c**) Schematic of an ommatidium with all major cell types labeled. (**d**) Eye cross-section illustrating how curvature affects field of vision and image resolution; ΔΦ is the interommatidial angle. (**e**) Developmental timeline of eye morphogenesis at 25°C, and major milestones; hAPF: hours after pupa formation. (**f**) 3D reconstruction of an ommatidium. Apical (left) and basal (right) views. 1°PC (light pink), 2°PC (pink), 3°PC, (yellow) and bristle cell complex (grey). PC: Pigment Cell. (**g** and **h**) Apical (g) and basal (h) confocal views through a 42 hAPF retina. Ommatidia form a hexagonal lattice apically (cyan hexagon). Orange rings mark the position of the grommet (h), the photoreceptors’ axon exit point, which in more apical sections aligns with the longitudinal axis of the ommatidium (g). Basally, elongated 2°PC profiles (pink) form a triangular mesh hinged at the grommets. The derived triangles (green) overlap the basal cell surfaces of the bristle cell complex (pseudocolored in grey), complementary to the 3°PC profiles (yellow). (**i**) View of a whole 42 hAPF retina with superimposed triangular mesh. Triangle size is color-coded (green-to-purple), revealing a dorsal/posterior to ventral/anterior gradient of increasing triangle size. A, anterior; P posterior. Scale bars: a, i = 100 µm; g, h = 10 µm.

The development of the compound eye is best understood in the fruitfly *Drosophila melanogaster*^5–7^. Cell differentiation and patterning of the retina into the mosaic of ommatidia starts in the late larval stage and continues after the larva begins its metamorphosis, in the early pupa. It is also during pupal life that the retina morphs into a 3D optical dome (Fig. 1e). This transformation occurs in three major steps. First, the thin pupal retinal epithelium becomes curved around 20 hours after pupa formation (hAPF), under the action of hydrostatic pressure which builds within the pupa ^8^. Then, at around 45–50 hAPF, ommatidial ancillary cells secrete the corneal lens, a hard polymer that coats the apical surface of the epithelium. Therefore, the final shape of the eye, including its local curvature anisotropies, is fixed by this time. Finally, starting at 55 hAPF, the retina thickens (up to 100 µm) as the ommatidial cells extend and their basal surfaces contract, finalizing the morphogenesis of a functional eye^8–11^. By the end of pupal development, the adult *Drosophila* eye shows a species-specific, stereotypic curvature (Fig. 1a,b). How this curvature is encoded in the fabric of the retinal tissue is not known.

### A patterned triangular mesh tiles the pupal retina

Early in pupal development, retinal cells acquire their position and remodel their morphology to shape the ommatidium as a 3D prism (Fig. 1f and Extended Data Fig. 1; ^9,12,13^). At this stage, apical (top) confocal views show the hexagonal lattice of ancillary interommatidial cells (IOCs) consisting of the secondary (2°PC) and tertiary pigment cells (3°PC), the four lens secreting cone cells and the sensory bristle cells complexes (Fig. 1g and Extended Data Fig. 1). Basal (bottom) confocal views show the cellular profiles of the IOCs coordinating their attachment around the afferent photoreceptor axons. In this organization, the IOCs form supracellular rings, called “grommets”, rich in extracellular matrix (ECM), which act as portholes through which the photoreceptor axons exit the retina^9,12,13^. Upon examining this basal surface, we observed a new level of organization: the elongated basal feet of the 2°PCs form triangles with the grommets as vertices, creating a continuous triangular mesh that spans the entire tissue (Fig. 1h).

Mechanical metamaterials are designed structures that consist of repetitive connected units. They are called “meta”materials because their unique mechanical properties come from how the units work together, not just from the material they are made of^4,14,15^. Combining physics engineering and computer science it has been possible to design *programmable* metamaterials, where the distribution of the units in 2D can control the 3D shape as loading is applied to the metamaterial^4,16^. One type of programmable metamaterials are bidimensional meshes in which the unit elements are triangles. In these “2D-triangular meshes” local curvature can be programmed by rationally varying the size of the triangles throughout the mesh^17–19^ (Box). The similarities between these metamaterials and the multicellular pattern of the retina led us to hypothesize that the developing compound eye might behave like a natural metamaterial, where the basal triangular mesh formed by the 2°PC would encode local curvature information. For this hypothesis to be true, three conditions must be met. First, the triangles of the mesh should be distributed across the 2D retina in a non-uniform, stereotyped pattern. Second, this 2D micropatterning should be instructive in generating the species-specific *Drosophila* 3D eye curvature. And third, perturbing the integrity of the triangular mesh should result in the eye losing its species-specific curvature.

#### Box. Generation and curvature quantification of 3D surfaces.

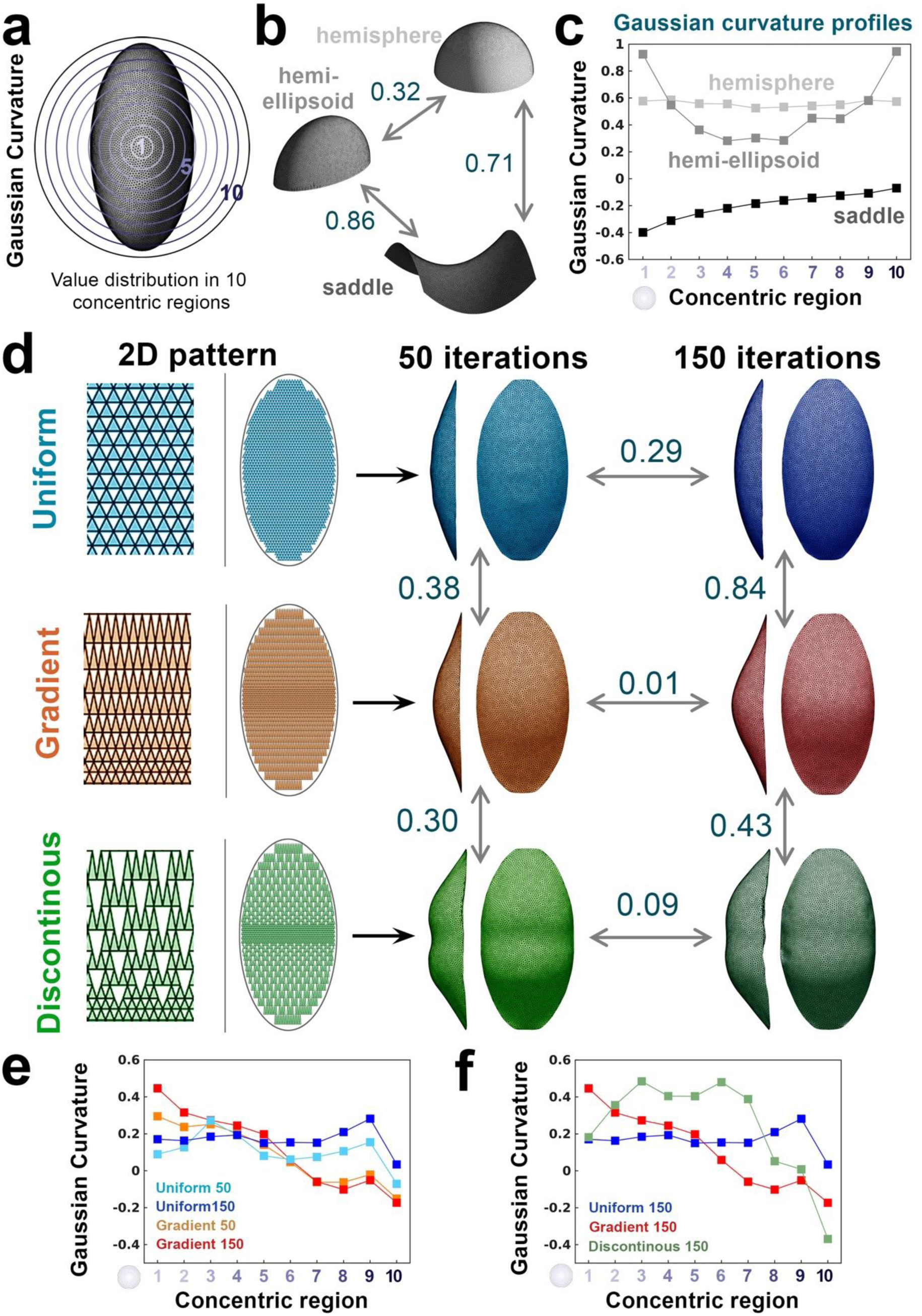

(**a**) To compare dome-like surfaces such as the *Drosophila* eye, it is necessary to evaluate the 3D curvature of the entire surface. Therefore, we calculated the Gaussian curvature distribution across the surfaces of interest. With this method, each curved surface is divided into ten concentric regions, and the integral of the dimensionless Gaussian curvature is calculated within each region (see Methods and Supplementary Methods). (**b**) As reference shapes, we use three idealized 3D surfaces: hemisphere, hemi-ellipsoid, and saddle. These illustrate typical curvature types: constant positive, varying positive, and negative curvature, respectively. The rigid perimeter is the same for all three patterns. (**c**) Each shape’s curvature is represented as a set of ten values (one per region), generating a *Gaussian curvature profile* (see Methods). The pairwise Gaussian metrics between the three reference shapes are shown in (b). The hemisphere and hemi-ellipsoid are similar (lower value), while the saddle is distinctly different from both. (**d**) To model curved metamaterial-like tissue formation, we apply pressure on a 2D triangular mesh enclosed in an elliptical rigid frame. We illustrate the differences in attained curvature using three patterns of triangle size: Uniform: identical triangle sizes (blue); Gradient: triangle sizes increase linearly from the equator toward the poles (orange); Discontinuous: same as the gradient but homogeneously removing 20% of the triangles in the upper bottom regions. Each 2D mesh is inflated over a series of computational iterations, and the resulting 3D shape is then processed for curvature analysis (Extended Data Fig. 3). The panel shows side views (yz and xz) of the processed surfaces generated by each pattern, at 50 (lighter colors) and 150 (darker colors) iterations. The xz views highlight overall shape and fine irregularities. (**e**) The Gaussian curvature profiles of the uniform (cyan/blue) and gradient (orange/red) surfaces are plotted after 50 and 150 iterations. The gradient pattern yields very similar 3D curvatures at both steps (Gaussian metric < 0.1). In contrast, the uniform pattern develops more noticeable curvature differences between 50 and 150 steps (Gaussian metric = 0.29), reflecting emerging irregularities during inflation. (**f**) Final Gaussian curvature profiles after 150 iterations for the three patterns showing poor correlation between them. Their pairwise Gaussian metric values are indicated in (d), reflecting the divergence of their 3D curvatures.

To analyze the pattern of triangle size of the basal mesh, we imaged and segmented the basal surface of pupal retinas before lens secretion. Meeting our first condition, we found the size of the triangles defined by the lattice of 2°PCs was distributed as a gradient of increasing size from dorsal/posterior to ventral/anterior across the retina (Fig. 1i, Extended Data Fig. 1 and Methods). Therefore, the mesh of triangles is micropatterned.

### The patterned triangular mesh encodes curvature

Next, to test whether the micropatterning of the 2D triangular mesh encodes the curvature of the *Drosophila* eye, we developed a physical model of this mesh. Using this model, we could program any distribution of triangle size within a given perimeter and simulate the resultant 3D curvature upon applying pressure to the mesh (Box and Supplementary Methods). To analyze and compare curvature between samples, we used a Gaussian curvature-based metric^20^ (Box, Supplementary Table 1 and Methods). Finally, as our goal was to compare computational and biological structures, we developed a computational pipeline to segment images of adult eyes, enabling precise measurement of local curvature (Fig. 2a,b, Extended Data Figs 2 and 3, Supplementary Table 2, Supplementary Video 1 and Methods).

**Fig. 2.**
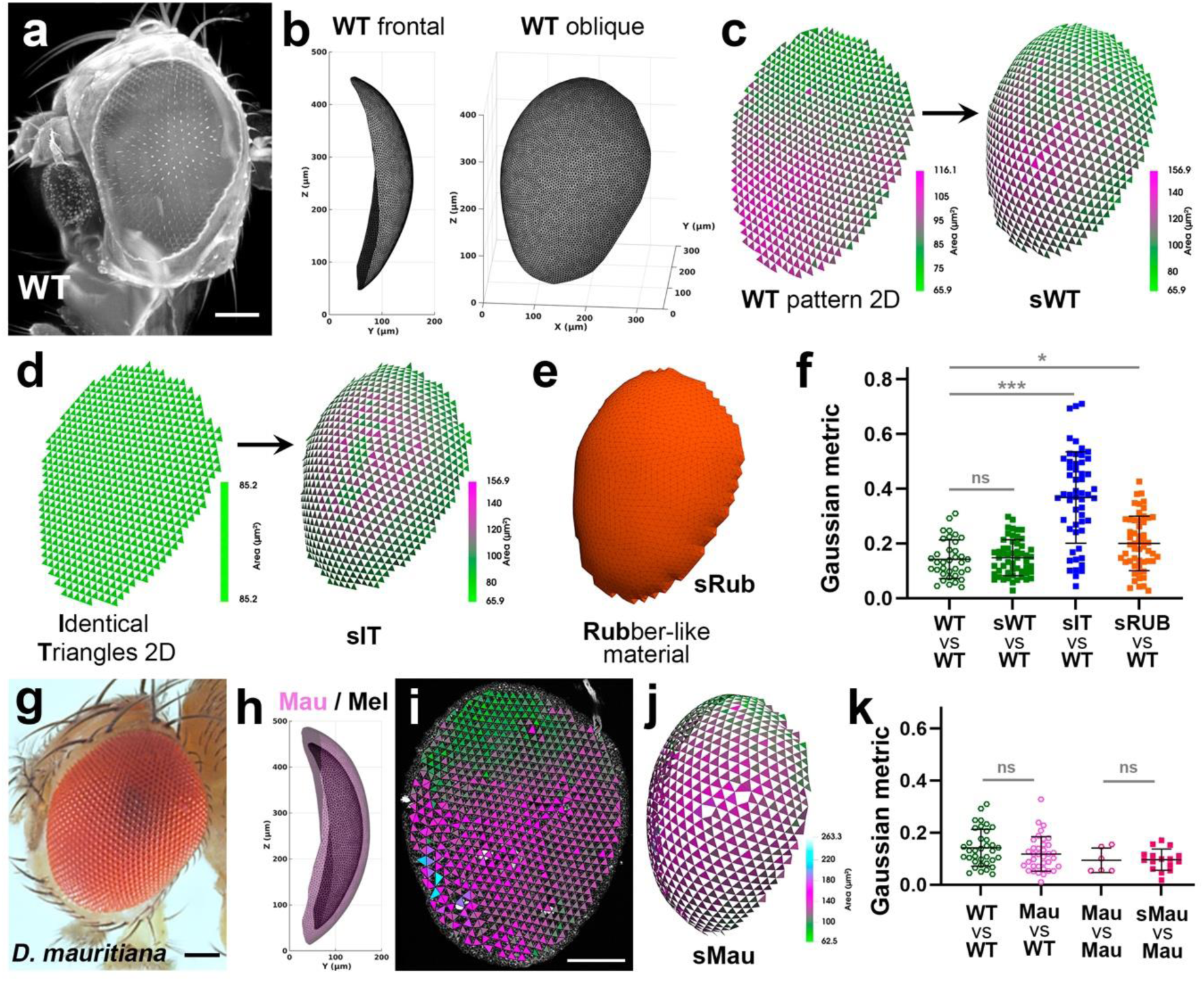
A physical model reproduces retinal curvature across *Drosophila* species. (**a**) Lateral view of a wild-type *D. melanogaster* head imaged using light-sheet microscopy. (**b**) Segmented eye surface (WT, frontal and oblique views) extracted from (a). (**c**) A patterned 2D triangle mesh extracted from a *D. melanogaster* pupal retina is used to generate a simulated 3D surface (sWT; see Methods). (**d**) For comparison, a 2D mesh consisting of triangles uniform in size, generates the sIT surface. (**e**) A third simulation (sRub) mimics a rubber-like material with the same perimeter. (**f**) Pairwise comparisons of the resulting 3D surfaces using the Gaussian metric (see Box); each dot represents one comparison. sWT is the only simulation that reproduces WT curvature. (**g**) Lateral view of an adult *D. mauritiana* head. (**h**) Frontal view of segmented *D. mauritiana* (Mau, pink) and *D. melanogaster* (Mel, grey) eyes, superimposed for comparison. (**i**) The *D. mauritiana* pupal retina displays a triangle size gradient similar to *D. melanogaster*. (**j**) sMau: the simulated 3D surface based on the *D. mauritiana* triangle pattern. (**k**) *D. melanogaster* (WT) and *D. mauritiana* (Mau) eyes are similar in curvature and sMau accurately reproduces the curvature of the adult *D. mauritiana* eye, showing a strong match with empirical data. Scale bars: a, g, i = 100 µm. Triangle size colored according to scale in (c), (d) and (i and j). Data shown as mean ± s.d. Gaussian metric distributions statistical tests: ns, not significant; *p<0.05; ***p<0.001.

To explore the link between triangular mesh and curvature, we programed three types of initial, 2D triangle patterns: (i) wild-type *Drosophila melanogaster* patterns derived from 42 hAPF retinas, “sWT”; (ii) uniform patterns with identical triangles, “sIT”; and (iii) continuous, rubber-like fine meshes composed of smaller triangles with scrambled orientations, “sRub” (see Methods for a detailed description; Fig. 2c-e, Extended Data Figs. 2 and 3). sRub was included to mimic the behavior of a homogeneous material^21^. To simulate the adult *Drosophila* eyes, we deployed each mesh within the perimeters measured from the pupal retinas, inflating each mesh until they best matched the mean depth of the wild-type eyes (Fig. 2b-e, Extended Data Figs. 2 and 3, Supplementary Video 2, Methods, Supplementary Tables 2 and 3). Finally, we computed the Gaussian metric in a pairwise manner (Supplementary Table 4). We found that local curvature of WT eyes was comparable across our samples (low values of Gaussian metric of “WT vs WT” in Fig. 2f, Extended Data Figs. 1 and 2, Supplementary Table 5, Methods). Remarkably, and validating our computational model, our analysis showed the curvature of the simulated WT (sWT) was very similar to the WT, while those sIT or sRub were significantly different (Fig. 2f, Supplementary Table 5, Methods). Therefore, only the triangle micropattern of *Drosophila* retinas was able to morph into the adult 3D eye curvature.

To further challenge our hypothesis, we tested whether there was a correlation between eye curvature and the pattern of triangles in the retinal mesh in other fly species. For this, we chose to examine the eye of *Drosophila mauritiana*, a species that diverged from *D. melanogaster* about 4 Myrs ago^22^ (Fig. 2g-k, Extended Data Fig. 4). *D. mauritiana* has larger eyes (Fig. 2g, Extended Data Fig. 4) as its retinas comprise more and larger ommatidia when compared to *D. melanogaster*^23^. The comparison of *D. mauritiana* (“Mau”) and *D. melanogaster* (“WT”, “Mel” in Fig. 2h) eyes showed that, despite their size difference, they have very similar curvature (Fig. 2k, Supplementary Tables 2 and 5). According to our hypothesis, the retina of *D. mauritiana* should present a patterned triangular mesh similar to that of *D. melanogaster*, a prediction we verified after analyzing the basal surface of *D. mauritiana* pupal retinas (Fig. 2i; compare with Fig. 1i). Moreover, incorporating the segmented triangle micropatterns into our computational model predicted the curvature of the *D. mauritiana* eye with great precision (Fig. 2k; see also Extended Data Fig. 4, Supplementary Table 5). These results further indicate that the 2D patterned triangular mesh encodes eye local curvature across *Drosophila* species.

### Disruption of the triangular mesh alters curvature of the compound eye

In our hypothesis, disruption of the 2D triangular mesh lining the basal surface of the retina should preclude curvature programming. As a consequence, the resulting eyes should lose their stereotypic curvature. To disrupt this mesh, we used RNAi (IR) to target the expression of Talin, a protein required for Integrin-mediated attachment of the 2°PCs to the grommet^12,13,24^ (genotype GMR-G4>talin_RNAi, “*talin-IR*”, see Methods). In this genotype, the 2°PCs lose their attachment to the grommets, which causes the disconnection of the mesh and affects the basal geometry of these cells (Fig. 3a and Extended Data Fig. 5). Despite these basal disruptions, apical patterning remains largely unaffected (Fig. 3b; and ^12,13,25^). While the *talin-IR* eyes were shaped as a dome-like structure like the WT (Fig. 3c and d, adult eye labelled as Talin in the figure), their curvature was markedly different from that of WT eyes (Fig. 3e-h, Supplementary Table 5). Notably, the disruption of the mesh produced large curvature variability (Fig. 3h, Supplementary Table 5), indicating that the triangular mesh is critical for the robustness of the 3D curvature. Breaking mesh connectivity should result in the retina losing the metamaterial properties –i.e *talin-IR* retinas should behave like a homogeneous material. To investigate this prediction, we simulated 3D curvature using the rubber material, described in Fig. 2e, which we framed within the perimeters of *talin-IR* retinas (“sTalin”, Extended Data Fig. 5 and Methods). This simulation led to a similar increase in variability to that we observed for adult Talin eyes (Fig. 3i and Supplementary Table 5). Altogether, our results support the concept that the metamaterial properties of the developing *Drosophila* retina, encoded in the patterned triangular mesh, are responsible for the reproducibility of the 3D curvature of the compound eyes.

**Fig. 3.**
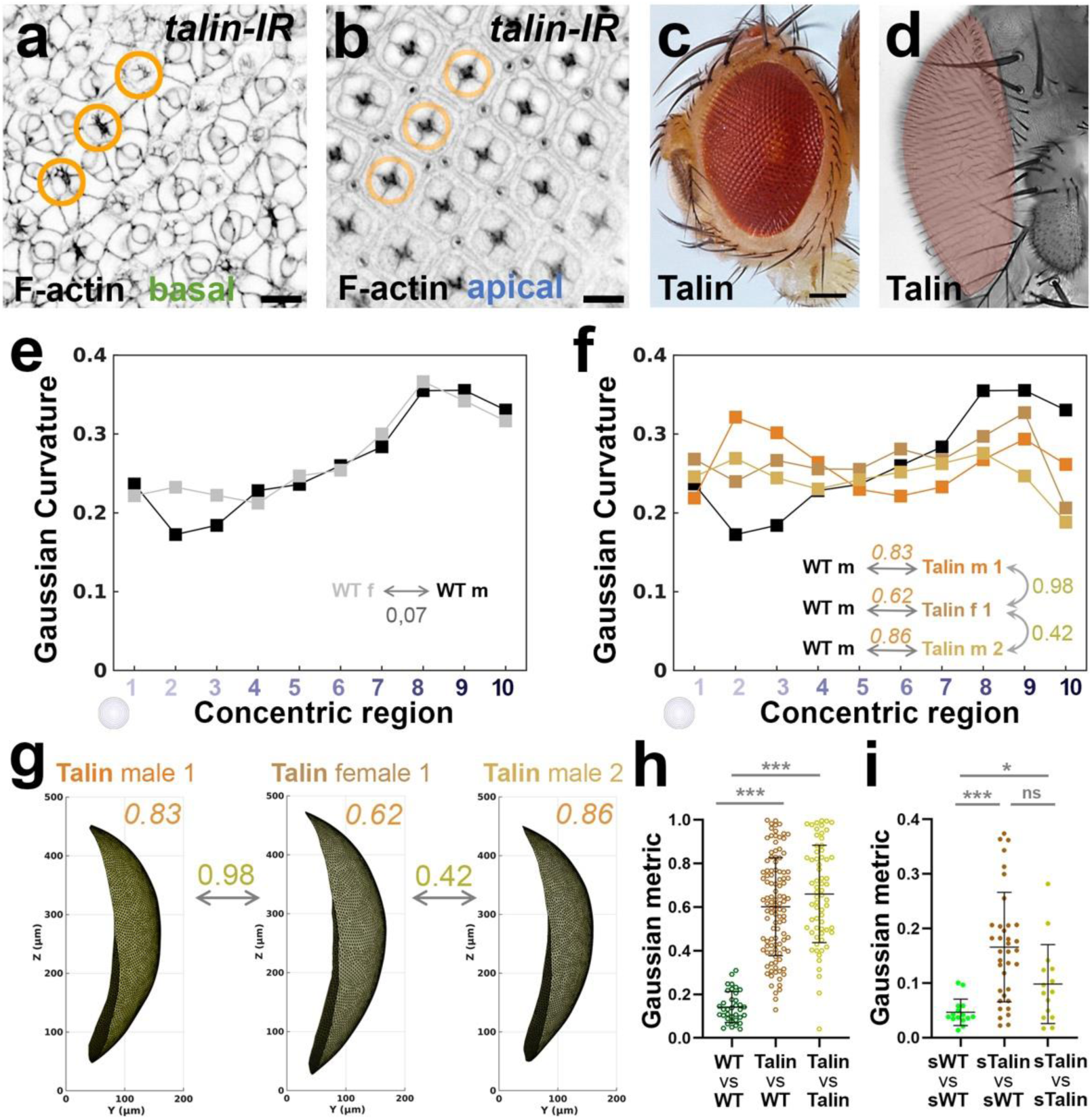
The patterned triangular mesh encodes for local curvature. (**a** and **b**) Confocal views through a *talin-IR* pupal retina stained with phalloidin to visualize the F-actin at the basal (a) and apical (b) surface of the retina. Orange ring marks the ommatidial central axis. (**c** and **d**) Lateral (c) and frontal (d) views of a *talin-IR* adult eye (“Talin” in the figure), showing that these eyes are curved and present minor alterations. (**e**) Reference Gaussian curvature profiles for a WT eye from a female (grey) and a male (black). Their similarity is indicated by a very low value of their pairwise Gaussian metric (0,07). (**f**) Combined dataset of Gaussian curvature profiles including a WT and three Talin surfaces, used to compute the pairwise Gaussian metric. Curvature of Talin eyes is consistently distinct from WT. (**g**) 3D reconstructions of three segmented Talin adult eyes. Bronze, italicized numbers indicate the Gaussian metric calculated from pairwise comparisons between each Talin surface and a wild-type (WT) male retina (see f). Gold numbers show the metric between each pair of Talin surfaces. (**h**) Distribution of Gaussian metrics for these comparisons, showing that Talin eyes differ significantly from WT and display higher variability. (**i**) Equivalent comparisons using computationally generated reference surfaces confirm these trends. Scale bars: a, b = 10 µm; c = 100 µm.

## Discussion

Since D’Arcy Thompson’s foundational work, it has been clear that understanding biological form requires examining the interplay between the principal forces and material properties at work during development^26^. During animal development, there are multiple examples where the physical forces that drive morphogenesis are spatiotemporally controlled at supracellular scales^27,28^. This idea is being pursued to engineer shapes in artificial tissues *in vitro*, through the programing of force asymmetries or microfabrication of 2D and 3D environments^29,30^. Here we have found that the compound eye uses a novel strategy to program a 3D shape *in vivo*: a uniform hydrostatic pressure acts on a patterned non-homogeneous tissue, the developing retina. This morphogenetic mechanism derives from two linked properties: the metamaterial quality of the tissue and the fact that it can be programmed genetically. The metamaterial behavior emerge from the specific mechanical coupling between 2°PCs, which is integral to the whole epithelium^11^. The connection of these cells through the grommets gives rise to the triangular mesh. The second part is the ability of locally controlling the size of the triangles, making it possible to establish a pattern responsible for the micrometer-scale, species-specific curvature anisotropies of the eye. A key question moving forward is which genetic mechanisms translate positional information into cell size regulation. The use of synthetic metamaterials with rationally-designed properties is fast expanding^31^, with new applications such as patches to give structural support to infarcted myocardium, vascular stents or wound dressings to aid skin healing^4^. To our knowledge, the developing retina of flies is the first instance of a natural metamaterial in which its properties are programmed genetically. By revealing how local geometry can be embedded in tissue architecture, this work introduces a novel strategy for the rational design of shape-programmable 3D biological surfaces, with potential implications extending from synthetic morphogenesis to clinical applications.

In addition to specifying the target morphology, a problem biological systems face is that of precision -reaching the species-specific morphology despite intrinsic and extrinsic noise. The need for precise curvature control has been made especially evident in studies of the *Drosophila* eye, where even subtle morphological defects can compromise optical function^32,33^. Moreover, local curvature variation must occur at the microscale, within a tissue only a few hundred microns across. Waddington proposed that phenotypic robustness should be the result of control mechanisms operating during development^34^. The phenomenon we describe here represents such a mechanism, where the robust and precise control derives from the metamaterial properties of the retina. From a design perspective, it is not clear why the triangular mesh is located at the basal surface rather than at the apical one. One possibility is that the basal surface is better suited for maintaining and guiding shape as it is directly exposed to the constant hydrostatic pressure, which during development has been shown to promote retinal curvature^8^. Once the apical lens is deposited and hardens, it likely serves as a permanent scaffold preserving retinal shape into adulthood. We also observed that the gradient in triangle size, from posterior/dorsal to anterior/ventral, mirrors a corresponding gradient in increasing lens size described in several *Drosophila* species, including *D. mauritiana* and *D. simulans*^35^. Since both curvature and lens diameter influence visual acuity, it is plausible that a control of ommatidial cell size co-regulates these two traits simultaneously. Considering the long evolutionary history of the compound eye, dating back to the Cambrian and coinciding with the explosive diversification of arthropods (reviewed in ^36^), it is tempting to speculate that mechanisms of curvature control and visual optimization, such as the one described here, may have played a role in the evolutionary success of insects and crustaceans.

## Supporting information

Extented Data Figures

## METHODS

### Fly strains & genetics

Flies were raised on standard food at 25°C. The following fly strains were used: *hth::YFP* (Kyoto:115109)^37,38^, *GMR-Gal4*^39^ (FlyBase: FBgn0020433; S0092-8674(00)81385-9 [pii]), *UAS-talin RNAi* (BDSC:33913)^40^, *D. mauritania* Tam-16 (gift from Alistair McGregor, Durham University, UK)^35^.

### Antibody staining and imaging

Retinas of appropriately staged animals were dissected in PBS on ice and fixed in 4% paraformaldehyde for 20 minutes at room temperature (RT). Retinas were washed in PBS-Triton 0.3% (PBS-T) then stained with primary antibody in PBS-T for 2hrs at RT or overnight at 4°C. Retinas were washed in PBS-T and then stained with secondary antibodies for 2h at RT or overnight at 4°C. Retinas were mounted in Vectashield (Vectorlabs)^41^. The following primary antibodies were used: Mouse N2 7A1 anti-Armadillo (1:200), mouse EXD B11M anti-Extradenticle (1:5) and rat DCAD2 anti-ECadherin (1:50). N2 7A1 Armadillo was deposited to the DSHB by Wieschaus, E. (DSHB Hybridoma Product N2 7A1 Armadillo)^42^. White, R. (DSHB Hybridoma Product EXD B11M), deposited EXD B11M to the DSHB^43^. DCAD2 was deposited to the DSHB by Uemura, T. (DSHB Hybridoma Product DCAD2)^44^. Anti-mouse or anti-rat secondary antibodies conjugated to CF 405S (Biotium, 20830) were used at 1:200 as appropriate, and ATTO 565 phalloidin (Sigma, 94072) was used at 0.4µM to visualize F-actin. Images of fixed retinas were acquired on a Zeiss 900 confocal microscope using the tile scan function.

### Preparation of adult heads and light sheet confocal imaging

This protocol, as the recipes used, are based on Susaki^45,46^. All incubations were performed at room temperature (RT) with agitation.

**Dissection**: Flies were euthanized in CO2 or on ice. They were then decapitated and the heads placed in a well containing 1X PBS. The proboscides were removed to allow further diffusion between the external medium and the interior of the head capsules. **Fixation**: The specimens were fixed in 4% paraformaldehyde in ethanol for 3-4 hours and then washed three times in pure ethanol for 1 hour each time. **Bleaching**: The heads were placed in tubes containing 10% H2O2 in ethanol until they were completely bleached (the time is variable and depends, for example, on whether the proboscis was fully removed or not). (Caution: This reaction produces oxygen. Leave the tube or well open during the first few hours of this step to allow the oxygen to escape. When the bubbling stops, the lid can be closed). This step may take from several days to 1,5 weeks for adult heads. Change the medium if it becomes pigmented. After completion of bleaching, wash 3 times with PBS 1x 1 h. **Clearing**: Heads were incubated in 50% Cubic-1/H2O for at least 3-6 hours to overnight. Then, incubated in Cubic-1 for 2 days. Next, incubated in Cubic-2/PBS for at least 3-6 hours to OV and then incubated in Cubic-2 for 2 days. Finally, they were incubated 3 times in glycerol/PBS 50%: first for 3-4 hours, second for OV, and third for 3-4 hours.

Microscopy was performed with a Zeiss Lightsheet 7 under a 5x objective. The heads were mounted on 1:1 glycerol/PBS columns with 1% w/v low melting agarose. The microscope chamber was filled with approximately 30 ml of 1:1 glycerol/PBS with a refractive index (RI) of 1.41. The software used was Zen Black (imaging) and Zen Blue (manual 3D reconstructions); the laser, 488 nm, which allows imaging of the cuticle autofluorescence.

### Statistical comparisons and interpretation

To evaluate the degree of similarity between simulations and adults (e.g., sWT and WT) we analyzed the statistical differences between the Gaussian metric distributions obtained from WT vs WT and sWT vs WT. In this example, both types of samples presented low values of Gaussian metric, so the absence of statistically significant difference was interpreted as indicating similar curvature between the two types of samples. We applied the same approach to compare the curvature of WT and *talin-IR* samples. In this case, the distributions of WT vs WT and Talin vs WT were significantly different, with the values for Talin being substantially larger. This result was interpreted as evidence of a difference in curvature and loss of robustness among the Talin samples.

The comparative analysis of the different Gaussian curvature profiles vector was conducted using a univariate statistical framework (Supplementary Tables 5 and 6). This approach enables the assessment of whether two datasets originate from populations with similar distributional properties. The protocol consisted of the following steps:

1. **Assessment of normality and homogeneity of variance**: For each pairwise comparison of Gaussian curvature distributions across genotypes, we first tested for normality using the Shapiro–Wilk test, and for homogeneity of variance using the two-sample F-test.
2. **Parametric testing with equal variances**: If both normality and equal variances were confirmed, we applied an unpaired *t*-test to assess differences in the means between the two groups. In case of multiple comparisons having one distribution as control, we applied a parametric ordinary one-way ANOVA test.
3. **Parametric testing with unequal variances**: In cases where the data were normally distributed but variances differed significantly, the two-tailed Welch’s *t*-test was employed as a more robust alternative. In case of multiple comparisons having one distribution as control, we applied a parametric ordinary one-way ANOVA test.
4. **Non-parametric testing**: When the assumption of normality was violated, we utilized the Wilcoxon rank-sum test (also known as the Mann–Whitney U test) to compare the medians of the two groups. In case of multiple comparisons having one distribution as control, we applied a Kruskal-Wallis test.

### Segmentation of confocal imaging data from 42hAPF retinas and generation of the triangular mesh

Confocal stacks stained with ATTO 565 phalloidin (F-actin) were used to segment grommets without prior knowledge of retina orientation. Segmentation began with MATLAB’s *Volume Segmenter* (R2021b), followed by a deep learning pipeline based on a 3D U-Net CNN trained on a manually segmented stack^47^. The resulting probability maps were binarized with custom MATLAB code and manually corrected. Samples requiring excessive manual editing were excluded. Grommet centroids were then extracted and used to construct alpha triangulations via the *alphaTriangulation* function.

To build the retina’s triangular mesh, we implemented a custom region-growing algorithm. Starting from a manually selected triangle over a bristle cell complex (*initial triangle*), the algorithm iteratively added non-adjacent triangles until completion of *valid triangles* set. Remaining triangles were classified as *inverse triangles*. Surface area–based color coding was applied to each triangle.

For *talin-RI* retinas (*GMR>talin_RNAi*) the segmentation procedure differed from that employed for the wild type *D. melanogaster* (“WT”) and *D*. *mauritania* (“Mau”), since grommets were largely absent, precluding neural network segmentation. These samples were fully segmented manually by delineating boundaries in *Volume Segmenter*. Boundary coordinates were extracted via custom scripts and used in downstream analyses.

### Building the different simulations

#### sWT

To make the any triangulation, in the case the one defined by the set of valid triangles of WT, sWT, compatible with the finite element method computational model employed in this study^48–50^ (Supplementary Methods), several preprocessing steps are required:

1. **Quadratic Triangulation**: We converted the initial triangulation into a quadratic triangulation using a custom MATLAB routine. Each triangle was redefined with six nodes, three at the vertices and three at edge points, yielding two matrices, a **coordinate matrix** (*coordinates*) of size and a **connectivity matrix** (*elements*) with nodes ordered clockwise^51^.
2. **Dirichlet Boundary Conditions**: We imposed these conditions by identifying the boundary nodes using MATLAB’s *boundary* functions and fixing all their **degrees of freedom** (dofs) to zero. These include three translational displacements along the *x*, *y*, *z* directions (dof = 1, 2, 3) and two rotational displacements about the *x* and *y* axes (dof = 4, 5). The conditions were stored in the *fixnodes* matrix to ensure model stability near boundaries.
3. **Nodal points**: The external forces were defined for internal (non-boundary) nodes, recorded in the *pointload* matrix. Initially, forces were applied in the *z*-direction, later updated during iterations to acto normal to each triangle’s surface (Supplementary Methods).

#### sWTi

To assess whether the gradient in the triangular pattern encodes the three-dimensional curvature of WT, we generated an inverted control pattern, referred to as sWTi, defined by the set of inverse triangles (Extended Data Fig. 2f,g). Once it was established, the same analytical and computational procedures were applied as used for sWT model.

#### sIT

For each of the sWTs, a corresponding uniform triangulation, sIT (Fig. 2d), was generated to create a mesh without spatial gradients in triangle size while preserving the original geometric pattern of valid triangles.

The process began by extracting the sWT boundary points and constructing a *uniform lattice* in the *x* − *y* plane. This lattice consisted of two alternating lines of points spaced by the average base *d_b_* and height *d*_ℎ_ of sWT triangles, producing congruent triangles of eqaul área across the domain. Points within the sWT boundary were selected and triangulated similarly tos WT methods, forming the sIT mesh.

Since the number of triangles in sIT could differ from sWT, an iterative adjustment of *d_b_* and *d*_ℎ_was performed until the number of triangles between sWT and sIT is reduced to within 5% (Supplementary Table 3).

#### sRub

For each triangulation based on the valid triangle set sWT, a corresponding rubber-like triangulation, sRub (Extended Data Fig. 2h), was generated to create a spatially continuous, non-structured and homogeneous mesh. Unlike the structured uniform triangulations (sIT), sRub was created using a simpler metho leveraging MATLAB’s Partial Differential Equation toolbox to automatically generate scrambled triangles within the sWT boundary. As a result, sRub meshes contained significantly more triangles than sWT, increasing computational demands. To balance simulation accuracy and efficiency, edge lengths were constrained between *H_max_* = 10 𝜇*m* and *H_min_* = 5 𝜇*m* (Supplementary Table 3).

#### sTalin

For the sTalin models, the same meshing procedure described for the sRub models was applied, including the specification of the maximum and minimum target edge lengths, *H_max_* and *H_min_*, respectively (Extended Data Fig. 5c). However, in this case, rather than utilizing the boundary of the corresponding sWT model, the triangulation was generated using the manually defined boundary of the *talin-RI* pupae.

#### sMau

The same procedure described for the sWT models was applied to Mau samples (Fig 2i).

#### sMau*

In the case of the sMau* models, it was necessary to refine the set of valid triangles of Mau samples due to the disruption of the characteristic triangulation pattern in specific regions of the anterior part of the retina (Extended Data Fig. 4f). This disruption resulted in the inclusion of computational triangles that did not accurately represent the basal surface of *Drosophila mauritiana* pupae. These non-representative triangles, that represent on average the 3% of the total triangles (Supplementary Table 3), were systematically identified based on morphological discrepancies with the expected basal surface geometry and subsequently excluded from the valid sMau triangulation set using a custom MATLAB script.

### 3D adult eyes image segmentation and postprocessing

To segment adult eyes independently from the heads across genotypes, we used the *Volume Segmenter* application with image data imported via a custom MATLAB script. Manual segmentation was performed followed by linear interpolation. The resulting label volumes were binarized and cropped for individual eye processing. A custom MATLAB algorithm extracted the apical surface by converting the segmented volume into a calibrated 3D point cloud. The point cloud was rotated so that the dorso-ventral axis lay in the *x* − *y* plane, aligning the apex (the point with the highest *z*-coordinate) normal with the *z*-axis. Spatial density was enhanced through interpolation with *griddata* function. The eye boundary was determined by projecting the point cloud onto the *x* − *y* plane and identifying its outer contour. Starting from the apex, the surface was propagated through connected points until the boundary was reached. Final boundary points were added and re-interpolated to yield a continuous apical surface.

Although the apical surface of the eye can be extracted from the segmented volume, the resulting representation is not suitable for the computation of dimensionless Gaussian curvature^20,52,53^.

To address these limitations, we developed dedicated MATLAB algorithm that consists of:

1. **Point Cloud Downsampling**: As proved in^54^ the dimensionless Gaussian curvature depends on the distance between a point and its neighbours. A downsampling strategy based on *farthest point sampling* (FPS)^55^ was implemented to iteratively select the subset of points that maximized spatial coverage while preserving the surface overall structure.
2. **Surface Smoothing:** To refine the mesh and reduce noise in the point cloud, we apply a smoothing operation based on Laplacian smoothing, which operates by iteratively adjusting the vertex positions to achieve a smoother surface while preserving the overall geometry. The smoothing process is performed as follows:

2.1 **Adjacency Matrix Construction**: At first, the adjacency matrix *A* ∈ ℝ*^N^*^×^*^N^* is built. Given the set of vertices *V* and faces *K* connectivity of the mesh (Supplementary Methods), for each triangular face {*i, j, k*} ∈ *K*, the adjacency matrix satisfies that *A*(*i, j*) = *A*(*j, i*) = 1.
2.2 **Laplacian Matrix Computation**: The Laplacian matrix *L* is computed using the relation *L* = *D* − *A*, where *D* is the degree matrix, defined as *D* =
2.3 *diag*(*A* ⋅ 𝟏), where 𝟏 is a column vector of ones.
2.4 **Iterative Smoothing**: The vertex positions are iteratively updated using the explicit smoothing scheme:

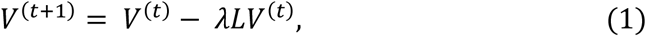

where 𝜆 is the smoothing factor, and 𝑡 denotes the iteration index. A smoothing factor of 0.1 was chosen and a total of 25 iterations are performed. This pair of values were chosen to balance convergence and preservation of geometric features, since small 𝜆 implies that each iteration produces modest changes in vertex positions, thereby avoiding excessive smoothing or distortion.
**Isotropic Explicit Remeshing**: Irregular meshes, can introduce significant local distortions in curvature computation^56^. To address this issue, an isotropic explicit remeshing step is implemented using *PyMeshLab*^57^. This remeshing algorithm is based on an energy minimization procedure and only the number of iterations is needed. We set the number of iterations to 25, which is sufficient to reach convergence of the remeshing energy minimum.

### Measuring dimensionless Gaussian curvature

The set of angles formed at vertex *v_i_*, by each of the incident triangles, *d_i_*, on it is denoted as 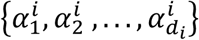, where each 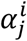represents the internal angle at *v_i_* within the *j*-the incident triangle. According to ^53,54,56^, the dimensionless Gaussian curvature *G_i_* over a neighborhood of vertex *v_i_* can be approximated by a discrete formulation:

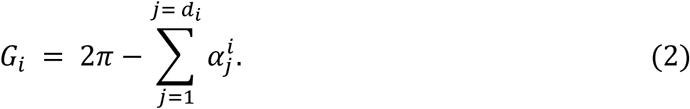

This formulation may yield misleading results due to inherent geometric and topological characteristics of the mesh such as open boundaries or topological defects. For vertices associated with such irregularities, defined as *degenerate vertices*, the computed values of *G_i_* does not reflect the curvature of the surface. These degenerate vertices are identified and removed using a custom-developed MATLAB algorithm:

1. **Detection of Boundary Edges and Vertices**: Edges that appear only once in the edge incidence map are classified as *boundary edges* and its vertices as *boundary vertices*. To improve curvature accuracy, all vertices in triangles containing any boundary vertex are also excluded from curvature calculations.
2. **Degeneracy Filtering Based on Geometric and Angular Criteria**: To ensure the integrity of curvature values, additional geometric criteria are enforced to identify degenerate vertices:

2.1 According to ^58,59^ vertices with any incident angle below 30° or above 90° may introduce a bias in *G_i_*. Although the isotropic explicit remeshing algorithm was applied, occasional violations still occur in regions near the boundary. To account for numerical accuracy, vertices with any incident angle below 33° or above 87° are identified as degenerate vertices.
2.2 In pupal samples, certain vertices located near the boundary may escape detection in the preceding filtering steps. Although such instances are infrequent, these vertices exhibit geometric characteristics analogous to true boundary points. For that reason, vertices for which |*G_i_*| > 0.01 are also identified as degenerate vertices.
3. **Identification of Interior Vertices**: Vertices not marked as boundary-related in the previous step are designated as interior vertices. These are the only candidates considered for reliable Gaussian curvature estimation.

### Shape similarity metric, the Gaussian metric

According to^20,60^ it is possible to define a similarity measure between two geometric models by means of a distance function *d*(*A, B*) that must satisfy the following properties:

● Non-negativity, *d*(*A*. *B*) ≥ 0.
● Symmetry, *d*(*A, B*) = *d*(*B, A*).
● Smaller values of *d*(*A, B*), more similar the shapes of *A* and *B*.

As demonstrated in ^20,61^, the integral of *G_i_* remains invariant under geometric transformations, so its distribution inherently defines a robust distance function for shape similarity.

As described in ^20^, the computation of that distribution is obtained by projecting the mesh vertices onto the *x* − *y* plane and transforming the point cloud, along *G_i_* values, into polar coordinates (𝑟*_i_,* 𝜃*_i_*), where the origin is defined as the geometric center of the point cloud (*x*_𝑐_*, y*_𝑐_) and the radial distance is computed as 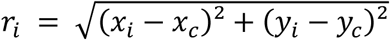.

The vertices are systematically ordered according to their radial distances 𝑟*_i_*. To partition the vertex set into *n* concentric regions, we define the *j*^𝑡ℎ^ region as comprising those vertices whose radial coordinates satisfy 𝑟*_i_* ∈ [𝑟*_j_,* 𝑟*_j_*_+1_), where 𝑟*_j_* denotes the 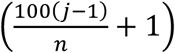 percentile of the distribution {𝑟_𝐼_}. This procedure yields *n* distinct groups of vertices, denoted as 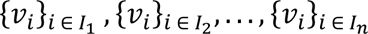, where the index set 𝐼*_j_* identifies all vertices belonging to the *j*^𝑡ℎ^ radial group. By construction, each group contains an equal number of vertices.

For each region 𝐼*_j_*, the integrated dimensionless Gaussian curvature 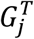, called *Gaussian curvature profile*, is computed as follows:

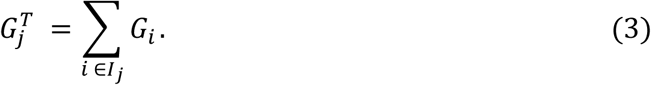

A vector 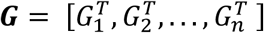 is constructed, that represents the surface. Consequently, the problem of comparing original surface meshes is reduced to the comparison of their 𝑮 vectors (Box). To this end, a distance function *d_G_* is defined as follows:

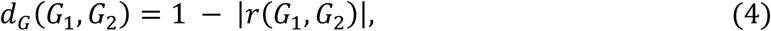

where *d_G_* ∈ [0,1) and |𝑟(*G*1*, G*2)| is the absolute value of Pearson correlation coefficient defined as:

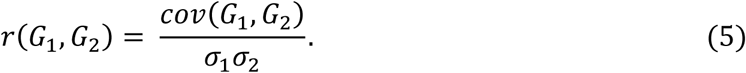

The distance function *d_G_* will henceforth be referred to as the *Gaussian metric*. Values near zero indicate high shape similarity between meshes, while values near one indicate significant shape dissimilarity (Box). According to ^20^, *n* ∈ [5,10], so we have adopted *n* = 10 to maximize the sensitivity of the Gaussian metric.

### Matching adult size and selecting the appropriate iteration

Following completion of all simulations generated by the computational model, 1000 for sWT, sWTi, sIT, sMau and sMau*, 2000 for sRub and sTalin, the resulting meshes were postprocessed using the same procedure described for adult specimens (Methods, 3D adult eyes image acquisition, segmentation and postprocessing). In this context, 75 iterations of the mesh smoothing algorithm were applied. This number was chosen to accommodate the fixed boundary conditions, which impose geometric constraints on curvature near the boundary and within regions that must remain free from negative curvature or irregular meshing artifacts. Given that the employed smoothing technique relies on the adjacency matrix, it enables selective regularization: boundary-adjacent regions are smoothed appropriately, while curvature in unaffected areas is preserved, thereby preventing excessive flattening due to oversmoothing.

Among the resulting iterations, it is necessary to select the one that best approximates the morphology of the adult specimens corresponding to the given genotype. This selection process is non-trivial, as the minimum of the Gaussian metric is not guaranteed to be unique, nor does it necessarily yield a configuration that accurately reflects adult morphology (Extended Data Fig. 3c). This limitation arises because the Gaussian metric, based on the distribution of dimensionless Gaussian curvature, does not account for absolute size.

To incorporate both shape and scale, ensuring that the selected iteration reflects not only the curvature distribution but also the physical size of the adult specimens, the following criteria were established to guide the selection of the optimal iteration:

1. **Size criterion**: To account for adult size, we first assessed whether significant differences existed between the lengths of the major axis in adults and pupae of the corresponding genotype (Supplementary Tables 2 and 3). As no statistically significant differences were observed in either case (Supplementary Table 6), the depth of the raw adults (preprocessing ones) was defined as the difference between the mean depth of the three-dimensional boundary and the depth of the deepest point. For each genotype, the acceptable depth range was defined as the mean adult depth ± one standard deviation. Any simulated specimen whose depth fell within this interval was considered to exhibit a depth-to-axis ratio statistically indistinguishable from that of the adults, thereby satisfying the size criterion (Extended Data Fig. 3b).
2. **Gaussian Metric Criterion**: For each raw iteration of the simulations (Extended Data Fig. 3a), depth was computed, as the maximum value of the z-coordinate, given that the boundary was constrained to z = 0. Additionally, G was calculated, along with < *d_G_* >, defined as the mean Gaussian metric from the distribution of adult individuals of the same genotype. This procedure yielded a pair (depth, < *d_G_* >) for each iteration. Among the iterations whose depths fell within the genotype-specific confidence interval defined by the size criterion, the one exhibiting the minimum Gaussian metric was selected (Extended Data Fig. 3c), thereby ensuring both geometric fidelity and curvature-based similarity to the adult morphology.

## Data availability

Code is available at: https://github.com/ComplexOrganizationOfLivingMatter/DrosophilaEyeCurvature. All other data are available in the main text or the supplementary information.

## ACKNOWLEDGMENTS

This work was supported by grants PID2022-137101NB-I00, AEI/10.13039/ 501100011033/FEDER UE (AEI/MICIN) (LME); PID2021-122671NB-I00 (AEI/MICIN) (FC); and PID2022-137101NB-I00 (AEI/MICIN) to JAA-SR. J.G.-G. was funded by a ‘Contrato predoctoral para la formación de doctores’ (PRE2020-093682) from the AEI/MICIN. Additional support from the E.U. COST action CA22153 ‘European Curvature and Biology Network’ (EuroCurvoBioNet) to LME, CEX2020-001088-M (AEI/MICIN) to FC, and LifeHUB research consortium (PIE-202120E047-ConexionesLife (CSIC)) to LME and FC is acknowledged. Work in the Pichaud lab is funded by grants from the MRC (MR/Y012089/1) and the BBSRC to FP and RW (BB/R00069). The *Drosophila* mauritiana Tam16 wild type strain was a gift from Alistair McGregor (Durham University, UK). Confocal Light Sheet imaging was carried out at ALMIA, CABD.

## AUTHOR CONTRIBUTIONS

LME, FC, and FP formulated the project. JG-G performed the computational design assisted by JAS-H. RW and JT performed the fly experiments, processed the samples and obtained the images. JG-G processed the images and performed the computational experiments. JAA-S assisted with image processing. JG-G, RW and JT analyzed the data. JG-G, RW, FP, FC, and LME wrote the paper with input from all authors.

