## Supplementary material for "A Natural Programmable Metamaterial Controls 3D Curvature of Compound Eyes": Suplementary Information

**SUPPLEMENTARY INFORMATION**

**Supplementary Methods**

### Mesh definition

According to [57] a mesh can be intuitively understood as a surface composed of flat, triangular elements joined together along their edges, forming a piecewise linear approximation of a shape. In our context, it is essential to distinguish between two aspects of a mesh: the arrangement of its elements and the spatial positions of its vertices. More precisely, a mesh $M$ can be defined as a pair $(K,V)$, where $K$ is a *simplicial complex* encoding the connectivity between vertices, edges, and faces, and thus characterizing the mesh's topology; and $V = \{v_1, \ldots, v_m\}$, $v_i \in \mathbb{R}^3$, is a set of coordinates specifying the geometric location of the vertices, thereby determining the actual shape of the mesh in $\mathbb{R}^3$, which makes up the geometric realization.

A simplicial complex $K$ is defined as a collection built from a finite set of vertices $\{1, \ldots, m\}$ and a family of non-empty subsets of these vertices, known as *simplices*. By definition, every individual vertex {i} must be included as a simplex, and any non-empty subset of a simplex in $K$ must also belong to $K$[57,62]. The simplices are categorized by dimension: 0-dimensional simplices $\{i\} \in K$ are referred to as vertices, 1-dimensional simplices $\{i, j\} \in K$ as edges, and 2-dimensional simplices $\{i, j, k\} \in K$ as triangular faces.

A mesh can be geometrically realized as a surface in $\mathbb{R}^3$ through the following construction. Starting from a given simplicial complex $K$, one forms its *topological realization* $|K|$ in $\mathbb{R}^m$ by associating each vertex $\{1, \ldots, m\}$ with a corresponding standard basis vector $\{e_1, \ldots, e_m\} \in \mathbb{R}^m$. For any simplex $s \in K$, the set $|s|$ represents the convex hull of its vertices in $\mathbb{R}^m$ and $|K| = \cup_{s \in K} |s|$. A linear mapping $\phi : \mathbb{R}^m \rightarrow \mathbb{R}^3$ is then defined to map the standard basis vectors $e_i \in \mathbb{R}^m$ to the actual positions of the corresponding vertices $v_i \in \mathbb{R}^3$, thereby embedding the abstract complex as a piecewise linear surface in three-dimensional space.

The *geometric realization* of the mesh $M$ is given by the image of $K$ under the map $\phi_V(|K|)$, where the notation $\phi_V$ highlights that the mapping is entirely determined by the set of vertex positions $V = \{v_1, \ldots, v_m\}$. The map $\phi_V$ is referred to as an *embedding* if it is injective, meaning that $\phi_V(|K|)$ contains no self-intersections. However, not all choices of vertex positions $V$ yield an embedding; only certain configurations of $V$ ensure that $\phi_V$ is one-to-one.

If $\phi_V$ is an embedding, then every point in the geometric realization $\phi_V(|K|)$ has a unique pre-image in $|K|$. This means that any point on the embedded surface can be parameterized by identifying its corresponding point in the abstract simplicial complex. The vector $\lambda$ such that $\phi_V(\lambda) = p$ where $p \in \phi_V(|K|)$ , is referred to as the *barycentric coordinate vector* $p$ with respect to $K$. These barycentric coordinates are convex combinations of the standard basis vectors in $\mathbb{R}^m$ corresponding to the vertices of a single face in $K$. Importantly, any barycentric coordinate vector has at most three non-zero entries: exactly three if the point lies in the interior of a triangular face, two if it lies on an edge, and one if it coincides with a vertex.

**Generation of box surfaces**

To simulate 3D metamaterial-like tissue formation, we apply a gradually increasing pressure to a 2D triangular mesh constrained within a rigid elliptical frame. The resulting differences in generated curvature are analyzed across three distinct triangle size distributions: Uniform, Gradient, and Discontinuous (Supplementary Table 1, Box). Each of these triangular patterns was generated as follows:

1. **Uniform pattern**: This pattern was generated using the same meshing procedure described for the sIT simulations. However, in this case, the total number of triangles was matched to that of the gradient pattern.

2. **Gradient pattern**: An elliptical boundary with a major-to-minor axis ratio of 0.5 was defined. Within this domain, a non-uniform triangular mesh was constructed using a method analogous to that employed for the sIT simulations, with key modifications. Specifically, instead of a uniform distribution, the mesh was designed to maintain a constant base triangle length $d_b$, while imposing a vertical gradient on the triangle height $d_h$. The height of each successive horizontal row of triangles was reduced by 5% relative to the previous one. This process continued until the height reached 20% of the initial value. From that point, a region of uniform triangle height was generated up to the midpoint of the ellipse along the $y$-axis. To preserve symmetry, the lower half of the pattern was constructed by mirroring this procedure across the horizontal axis.

3. **Discontinuous pattern**: This configuration was derived from the gradient pattern by homogeneously removing 20% of the triangles from both the upper and lower regions of the mesh.

**Farthest point sampling**

This method iteratively selects a subset of points that maximizes spatial coverage of the surface while preserving its overall structure. Starting from an arbitrary initial point, each subsequent point is chosen to be the one that lies farthest, in Euclidean distance, from the nearest previously selected point. Formally, given a point cloud $P = \{p_1, p_2, \ldots, p_N\} \subset \mathbb{R}^3$, and a desired number of sampled points $n < N$, the algorithm constructs an index set $I \subset \{1, \ldots, N\}$ of cardinality $n$ such that the minimum distance between any unselected point and the selected set is maximized at each iteration. The result is a sparser, yet well-distributed, point cloud.

**Computational model to deploy the triangular mesh**

A finite element method (FEM)[50], was implemented in MATLAB to simulate the mechanical response of thin or thick shell structures using a triangular shell element. The model is based on a 6-node quadratic triangular shell element, capable of capturing both membrane and bending behaviors. The formulation is suitable for geometrically nonlinear analyses and includes options for viscous stabilization, prescribed boundary conditions, and iterative load updates in the direction of nodal normals.

The mechanical behavior of the shell is governed by the principles of linear elasticity. The shell kinematics are described using Mindlin–Reissner theory, which accommodates transverse shear deformation, making the formulation applicable to both thin and moderately thick shells.

Each node of the triangular element possesses five degrees of freedom:

- Three translational components: $\boldsymbol{u_x}, \boldsymbol{u_y}, \boldsymbol{u_m}$.
- Two rotational components: $\boldsymbol{\theta_x}, \boldsymbol{\theta_y}$.

In our model, we assume that connected triangles can freely rotate at connection points between triangles thereby decoupling individual triangle rotations. To computationally implement this condition, we define duplicated nodes at connection points which are after kinematically linked across their translational components. This kinematic condition is imposed in our system through a penalty analogous to the imposition of prescribed displacements/rations in FEM[50].

The generalized strain vector at a Gauss point is divided into:

1. Membrane strains ($\boldsymbol{\varepsilon_m}$) — in-plane normal and shear strains.
2. Bending curvatures ($\boldsymbol{\kappa_b}$) — second derivatives of the displacement field in the mid-surface.
3. Shear strains ($\boldsymbol{\gamma_s}$) — due to transverse shear deformation.

Each component is related to the nodal displacements $u$ by:

$$\boldsymbol{\varepsilon_m} = B_m\boldsymbol{u}, \qquad \boldsymbol{\kappa_b} = B_b\boldsymbol{u}, \qquad \boldsymbol{\gamma_s} = B_s\boldsymbol{u}, \tag{6}$$

where:

- $B_m$: It is the membrane strain matrix. This matrix maps the translational degrees of freedom in the element mid-surface to the in-plane strains. Assuming a linear or quadratic interpolation of displacements:

$$B_m = \begin{pmatrix} \frac{\partial N_1}{\partial x} & 0 & 0 & \dots & \frac{\partial N_n}{\partial x} & 0 & 0 \\ 0 & \frac{\partial N_1}{\partial y} & 0 & \dots & 0 & \frac{\partial N_n}{\partial y} & 0 \\ \frac{\partial N_1}{\partial y} & \frac{\partial N_1}{\partial x} & 0 & \dots & \frac{\partial N_n}{\partial y} & \frac{\partial N_n}{\partial x} & 0 \end{pmatrix}.$$

  where $N_i$ are the shape functions for each node in the element.

- $B_b$: It is the bending strain matrix. This matrix relates the rotations, usually associated with out-of-plane curvature, to bending strains:

$$B_b = \begin{pmatrix} \frac{\partial \theta_{x,1}}{\partial x} & \frac{\partial \theta_{y,1}}{\partial x} & \cdots & \frac{\partial \theta_{x,n}}{\partial x} & \frac{\partial \theta_{y,n}}{\partial x} \\ \frac{\partial \theta_{x,1}}{\partial y} & \frac{\partial \theta_{y,1}}{\partial y} & \cdots & \frac{\partial \theta_{x,n}}{\partial y} & \frac{\partial \theta_{y,n}}{\partial y} \\ \frac{\partial \theta_{x,1}}{\partial y} + \frac{\partial \theta_{y,1}}{\partial x} & \frac{\partial \theta_{y,1}}{\partial x} + \frac{\partial \theta_{x,1}}{\partial y} & \cdots & \frac{\partial \theta_{x,n}}{\partial y} + \frac{\partial \theta_{y,n}}{\partial x} & \frac{\partial \theta_{y,n}}{\partial x} + \frac{\partial \theta_{x,n}}{\partial x} \end{pmatrix}.$$

  where $\boldsymbol{\theta_x}, \boldsymbol{\theta_y}$ are rotations about the global $x$ and $y$ axes, respectively.

- $B_s$: It is the shear strain matrix. This matrix captures the transverse shear deformation and is often simplified via reduced integration to avoid shear locking:

$$B_s = \begin{pmatrix} \frac{\partial N_1}{\partial x} & 0 & -N_1 & \cdots & \frac{\partial N_n}{\partial x} & 0 & -N_n \\ 0 & \frac{\partial N_1}{\partial y} & -N_1 & \cdots & 0 & \frac{\partial N_n}{\partial y} & -N_n \end{pmatrix}.$$

The total stiffness matrix for each element $K_e$ is constructed as:

$$K_e = \int_A (B_m^T D_m B_m + B_b^T D_b B_b + B_s^T D_S B_S)\, dA, \tag{7}$$

where:

- $B_m, B_b, B_s$: Strain-displacement matrices for membrane, bending, and shear components, respectively.

- $D_m, D_d, D_s$: Constitutive matrices for each component (see next section).

- $A$: Element area.

The integrals are evaluated using one-point reduced integration (three-point quadrature for triangles).

Constitutive material model

The material is modeled as isotropic and linearly elastic. Three constitutive matrices are used for the three physical mechanisms in the shell:

- Membrane forces:

$$D_m = \frac{Et}{1-\nu^2}\begin{pmatrix} 1 & \nu & 0 \\ \nu & 1 & 0 \\ 0 & 0 & \frac{1-\nu}{2} \end{pmatrix}$$

- Bending moments:

$$D_b = \frac{Et^3}{12(1-\nu^2)}\begin{pmatrix} 1 & \nu & 0 \\ \nu & 1 & 0 \\ 0 & 0 & \frac{1-\nu}{2} \end{pmatrix}$$

- Shear forces:

$$D_s = \frac{5}{6} G t I_2, \qquad G = \frac{E}{2(1+\nu)},$$

where $E$ is Young's modulus, $\nu$ is Poisson's ratio, $t$ is the shell thickness, and $I_2$ is the $2x2$ identity matrix. In the present work, $E/p = 6{,}825 \cdot 10^4$, $\nu = 3 \cdot 10^{-1}$ and $t = 1{,}25 \cdot 10^{-1}\ \mu m$. The chosen value of $E/p$ is given in a dimensionless fashion to ensure that for a selected applied pressure $p = 10^5$, results in consistent deformations while maintaining numerical stability in the simulation. The thickness $t$ was selected such that the ratio $t/h$, being $h$ the height of the triangle, remains within the bounds required by thin shell theory (Supplementary Table 3). These parameters are therefore chosen to satisfy the theoretical constraints of thin shell kinematics[63] while promoting a regularized and robust computational framework.

<u>Solution strategy</u>

The nonlinear equilibrium of the shell structure is solved through a quasi-static iterative procedure implemented using an updated Lagrangian scheme. Rather than relying on a residual-based convergence criterion, the algorithm executes a predefined number of iterations per load step, thereby enforcing consistency through gradual pressure application and geometric updates. This approach is particularly effective for thin-to-moderately thick shells where load-induced geometric nonlinearities must be captured, but material behavior remains linear.

At each load step, the global displacement vector $\boldsymbol{u}$ is initialized and incrementally updated across a fixed number of iterations. Subsequently, provided the updated displacement vector, mesh coordinates are appropriately updated. Then, at each iteration, the following steps are performed:

1. **Normal Directions Update and Load Update Procedure**: The local normal vectors at the surface nodes are recalculated using an area-weighted average of the normals

from adjacent triangular elements. This procedure ensures that externally applied surface loads remain correctly oriented relative to the evolving geometry. Once the face normals have been updated, the external load vector is recalculated to ensure that the applied forces remain aligned with the deformed surface. In that way, pressure is always being applied normally over the surface.

2. **Global System Assembly**: The global system assembly forms the structural core of the finite element method, wherein the element-wise stiffness matrices and load vectors are constructed and assembled into global matrices. The total stiffness matrix of the structure is obtained by looping over all elements and computing their contributions using numerical integration.

2.1. Element Geometry and Local System Transformation: For each element $e \in \{1, \ldots, n_{elem}\}$, the nodal coordinates are extracted using the connectivity matrix $elements$, which indexes into the nodal coordinate array $coordinates$. A local coordinate system $T_e \in \mathbb{R}^{3\times3}$ is constructed based on three selected nodes to define the element's mid-plane and local axes. This transformation matrix is used to express local quantities in a reference frame, called local system, aligned with the mid-surface of the shell:

$$c_{local} = c_{global} T_e^T. \quad (8)$$

2.2. Numerical Integration and Element Stiffness Matrix: Numerical integration is performed over each element using three Gauss points. At each Gauss point $g$, the strain-displacement matrices are computed and then, the corresponding contribution to the element stiffness matrix $K_e$ are integrated:

$$\begin{aligned} K_b^{(g)} &= B_b^T D_b B_b a_g w_g \\ K_m^{(g)} &= B_m^T D_m B_m a_g w_g. \\ K_s^{(g)} &= B_s^T D_s B_s a_g w_g \end{aligned} \quad (9)$$

The total element stiffness matrix is updated via:

$$K_e = K_e + K_b^{(g)} + K_m^{(g)} + K_s^{(g)}, \quad (10)$$

where $a_g$ is the area of the Gauss point and $w_g$ is the integration weight. This equation is the discretized version of equation (7).

2.3. Distributed Load Vector: Simultaneously, the distributed loading $p$ applied normal to the shell mid-surface is incorporated into the element force vector. The load vector contribution at Gauss point $g$ is given by:

$$\boldsymbol{f}_{\boldsymbol{e}}^{(\boldsymbol{g})} = N^T(p\boldsymbol{n} - \rho h g)a_g w_g, \tag{11}$$

where $N$ is the shape function matrix, $\boldsymbol{n}$ is the local normal vector (third row of $T_e$), $\rho$ is the shell density, $h$ is the shell thickness and $g$ is the gravitational acceleration vector. For the sake of simplicity, in the present work $\rho = 0$.

2.4. Assembly into Global Matrices: For each element, the degrees of freedom are mapped to global indices using the connectivity matrix and degree-of-freedom per node. The stiffness matrix $K_e$ and force vector $\boldsymbol{f_e}$ are assembled into the global stiffness matrix $K \in \mathbb{R}^{n_{dof} \times n_{dof}}$ and the global force vector. This step completes the discretization of the equilibrium equations over the entire domain.

3. **Viscous Regularization**: During the numerical simulations using the shell finite element model, it was observed that the system exhibits instabilities consistent with rigid-body motion, particularly uncontrolled rotational displacements of the triangular shell elements, as elements are freely allowed to rotate being these rotations uncoupled between elements. This behavior indicates that the linear system to solve is poorly conditioned due to a lack of sufficient constraints to suppress such non-physical motions.
To address this, a viscous regularization strategy is introduced. This approach conceptually embeds the shell in a fictitious viscous medium, such that the elements experience a damping force proportional to their velocity. This artificial viscous force has only a numerical meaning and it does not alter the solution significantly in the steady state but provides numerical stability by damping out rigid-body motions during the iterative solution process. The regularized system is reformulated as:

$$\left(\frac{c}{\Delta t} + K\right)\boldsymbol{u_{t+1}} = \boldsymbol{f_{t+1}} + \frac{c}{\Delta t}\boldsymbol{u_t}, \tag{12}$$

where $\boldsymbol{u_{t+1}}$ is the current displacement vector, $\boldsymbol{u_t}$ is the displacement vector from the previous iteration, $K$ is the stiffness matrix, $\boldsymbol{f_{t+1}}$ is the updated force vector and $c$ is the viscous damping matrix, serving as a regularization term, defined as:

$$c = \alpha I_{disp}, \tag{13}$$

where $\alpha$ is a small scalar regularization parameter and $I_{disp}$ is a diagonal matrix of size $n_{dof} \times n_{dof}$ with ones in positions corresponding to translational degrees of freedom and zeros elsewhere. The scalar coefficient $\alpha$ is calibrated to ensure the damping remains a small perturbation to the physical problem. It is computed as:

$$\alpha = \left(\frac{\max(K) + \min(K)}{2}\right) x\ 10^{-6} \tag{14}$$

This empirical scaling ensures that the regularization is effective enough to suppress instabilities without significantly altering the solution of the mechanical system. By incorporating this viscous regularization, the simulation gains robustness and numerical coherence, while preserving the validity of the thin shell formulation. The method serves as a non-intrusive stabilization mechanism for large deformation analyses where rigid-body motions may otherwise persist.
