## Supplementary material for "A Natural Programmable Metamaterial Controls 3D Curvature of Compound Eyes": Extented Data Figures

### EXTENDED DATA FIGURES

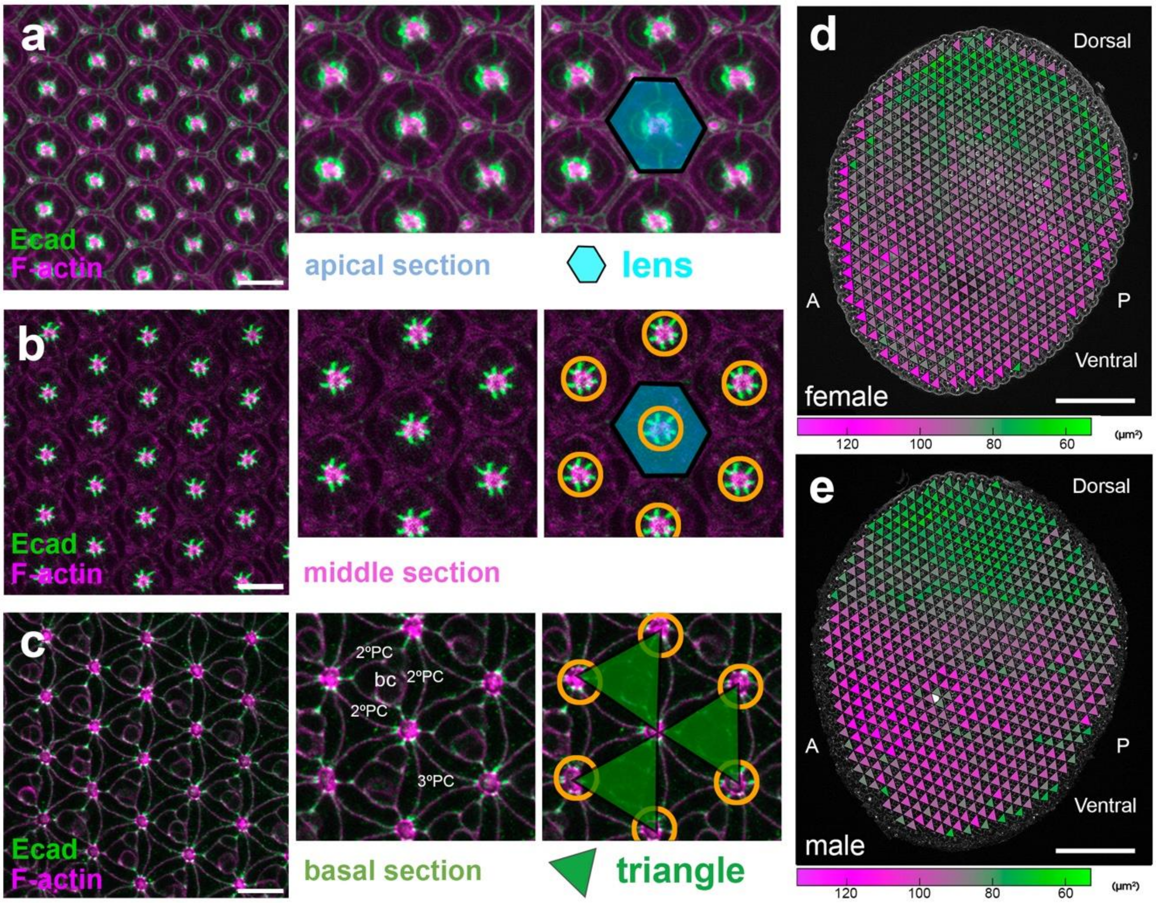


**Extended Data Fig. 1: A supracellular triangular pattern in the *Drosophila* pupal retina**. (**a** to **c**) Apical (a), middle (b) and basal (c) confocal views through the retinal epithelium at 42h APF. The panels show the correspondence of structures along the apicobasal axis of the ommatidium, from the apical hexagonal packing to the basal triangles formed by the feet of the secondary pigment cells. The grommets (orange circumference) are in the center of the hexagons and at the vertex of the triangles. Green: E-cadherin. Magenta: F-actin. bc: bristle cell complex; 2ºPC: secondary pigmentary cell; 3ºPC: tertiary pigment cell. (**d** and **e**) The dorsal (top) to ventral (bottom) gradient of increasing triangle size in female (d) and male (e) 42 hAPF retinas. View with superimposed triangular mesh. Triangle size scale is color-coded (green-to-purple). A, anterior; P posterior. Scale bars: a, b, c = 10 µm; d, e = 100 µm.

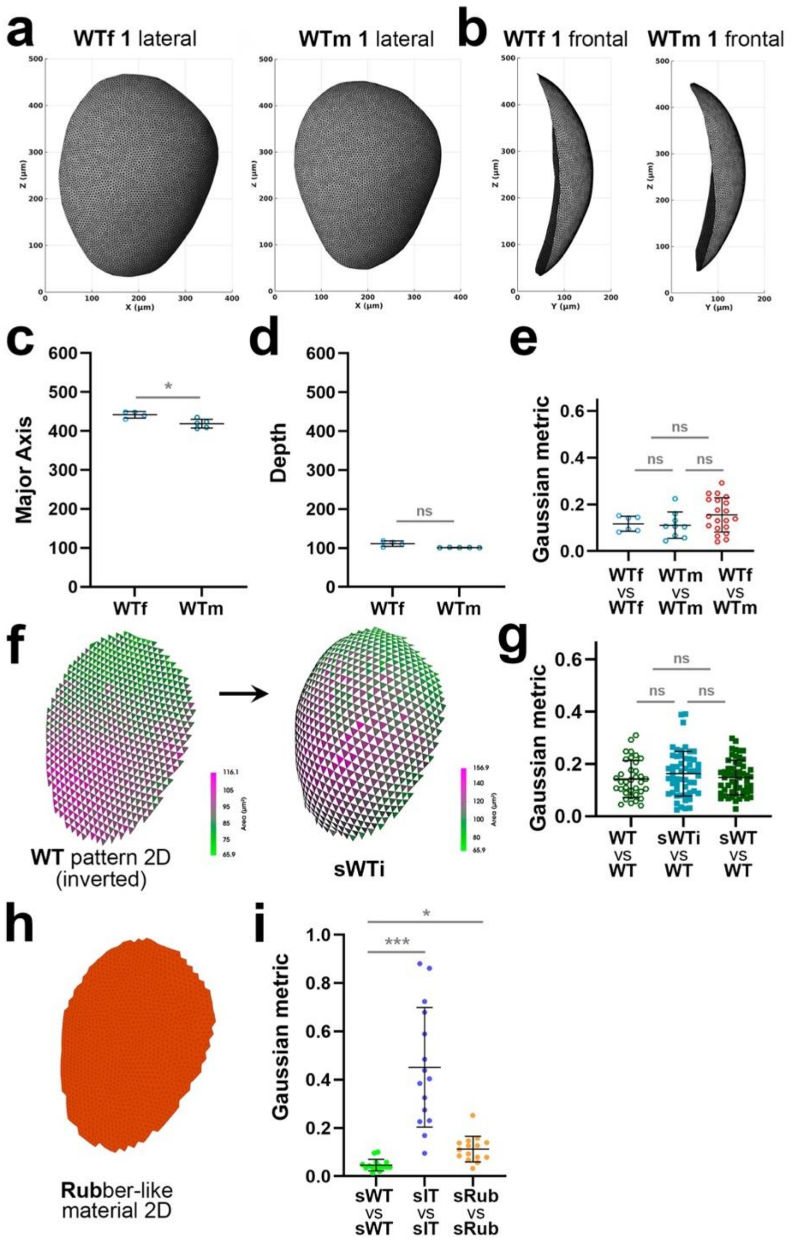


**Extended Data Fig. 2: Curvature comparisons of adult eyes and different patterns simulated computationally.** (**a** and **b**) Representative segmented eye surface (frontal (a) and lateral (b) views) of female and male flies ("WTf 1" and "WTm 1", respectively). (**c** and **d**) The adult female eye has a bigger major axis (c), but males and females present similar eye depth (D) (Supplementary Tables 2 and 6). (**e**) Distribution of Gaussian metrics for male and female comparisons, showing that they all present similar values with low variability. (**f**) A 2D triangle size map of the *D. melanogaster* pupal retina complementary to the one shown in Fig. 1c, but with the same outline, is used to generate a simulated 3D surface (sWTc; see Methods). Triangle size scale is color-coded. (**g**) Pairwise comparisons of the resulting WT, sWTc and sWT surfaces showing the high similarity between all of them. (**h**) A disordered pattern of small triangles is the initial 2D surface for the rubber-like material simulations. (**i**) Pairwise comparison showing that the WT patterns produce very similar curvatures that cannot be reproduced neither with the Identical Triangles ("sIT") nor the Rubber-like material ("sRub").

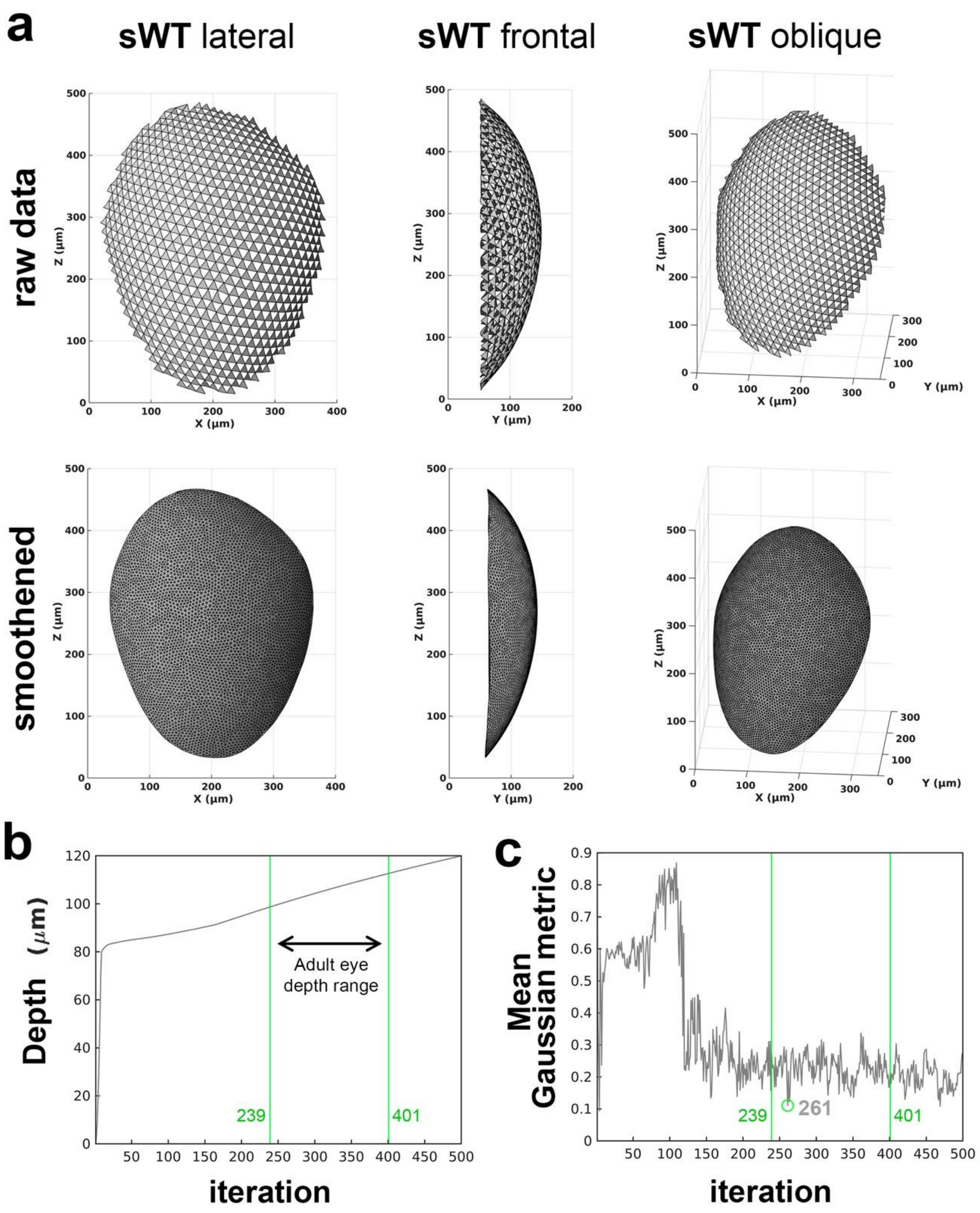


**Extended Data Fig. 3: Methodology for the comparison of adult eyes and models.** (**a**) Representative lateral, frontal and oblique views of the raw sWT (top row) and the corresponding smoothed surface that can be compared with the adult eyes (bottom row, see Methods). (**b**) Evolution of raw sWT depth (grey) as a function of iterations. The mean ± s.d of WT depth defines the selection window for identifying the best-fitting sWT iteration (green lines: iteration 239-401). (**c**) Evolution of mean Gaussian metric between sWT and WT (grey) across iterations. Within the selection window (green lines: iteration 239-401) the selected iteration that corresponds to sWT is identified as the one with the lowest WT mean Gaussian metric.

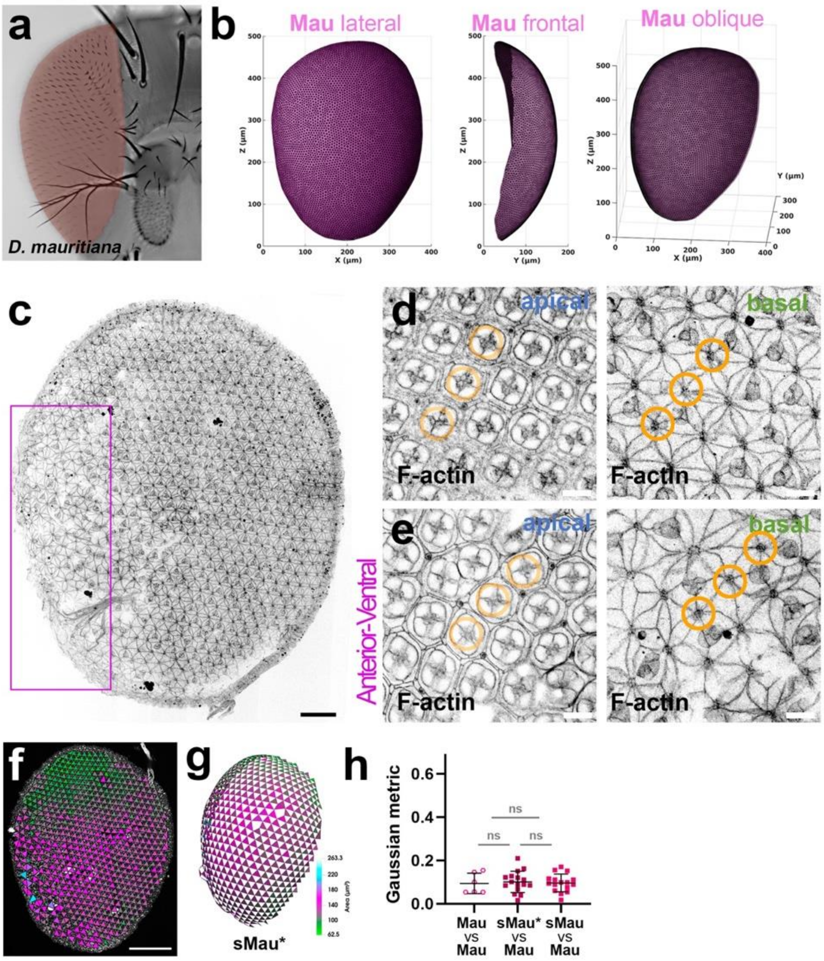


**Extended Data Fig. 4**: **Analysis of Curvature in the *Drosophila mauritiana* Eye.** (**a**) Frontal view of an adult D. mauritiana eye. (**b**) Lateral, frontal, and oblique views of a representative segmented eye. (**c**) Tile-scan confocal image of a full retina stained with phalloidin to visualize the F-actin at the basal surface. The boxed region shows mild defects in the basal triangular pattern of 2º PCs restricted to this region. (**d**) Apical and basal confocal views from the posterior part of the retina. The apical packing is normal. In the basal view, the triangular mesh pattern is visible despite the absence of some bristle cell complexes. (**e**) Apical and basal confocal views from the boxed region in (c). The pattern in the basal region is disorganized, and some of the triangle "sides", as formed by the 2ºPC basal feet, are missing. (**f**) *D. mauritiana* pupal retina from Fig. 2i where some triangles have been removed from places where 2ºPC basal feet are missing. (**g**) sMau*: the simulated 3D surface based on the *D. mauritiana* triangle pattern with triangles removed. Triangle size scale is color-coded for (f and g). (**h**) Distribution of Gaussian metric of *D. mauritiana* eyes (“Mau”), sMau and sMau*. No curvature differences were detected in any of the comparisons. The similarity of the metrics between sMau (with a full complement of triangular segmentation based on grommet as triangle vertices) and sMau* (with some triangles removed to mimic the “defects” observed, d basal), indicates that the small number of missing triangles in the anterior/ventral region of Mau retinas does not significantly affect curvature (Supplementary Table 5). Scale bars: c, f = 100 µm; d, e = 10 µm.

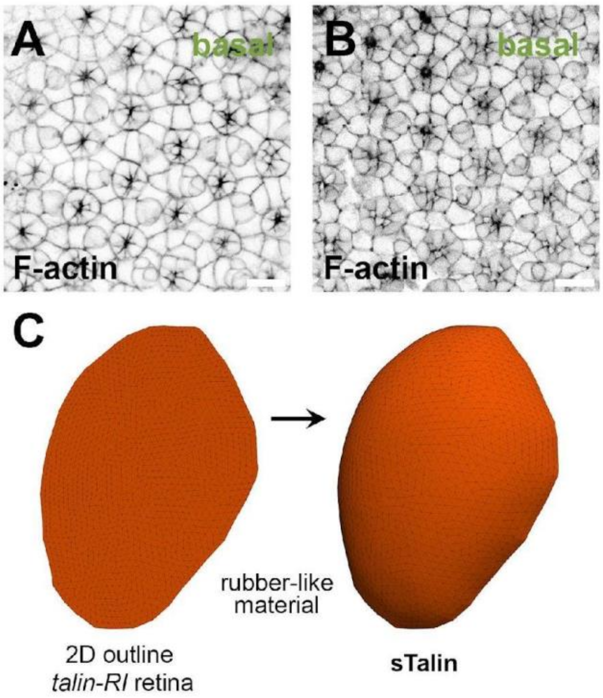


**Extended Data Fig. 5: *talin-IR* phenotype and its simulation.** (**a** and **b**) Representative basal confocal views of two different *talin-IR* pupal retinas. (**c**) A continuous fine mesh composed by triangles with scrambled orientation inside of the perimeter of the *talin-IR* pupal retina is the initial 2D surface that is inflated to produce the sTalin simulation (see Methods).
